# FITTING OF HYPERELASTIC CONSTITUTIVE MODELS IN DIFFERENT SHEEP HEART REGIONS BASED ON BIAXIAL MECHANICAL PROPERTIES

**DOI:** 10.1101/2021.10.28.466240

**Authors:** Fulufhelo Nemavhola, Thanyani Pandelani, Harry Ngwangwa

## Abstract

Heart failure remains one of the leading causes of death especially among people over the age of 60 years worldwide. To develop effective therapy and suitable replacement materials for the heart muscle it is necessary to understand its biomechanical behaviour under load. This paper investigates the passive mechanical response of the sheep myocardia excised from three different regions of the heart. Due to the relatively higher cost and huge ethical demands in acquisition and testing of real animal heart models, this paper evaluates the fitting performances of five different constitutive models on the myocardial tissue responses. Ten sheep were sacrificed, and their hearts excised and transported within 3h to the testing biomechanical laboratory. The upper sections of the hearts above the short axes were carefully dissected out. Tissues were dissected from the mid-sections of the left ventricle, mid-wall and right ventricle for each heart. The epicardia and endocardia were then carefully sliced off each tissue to leave the myocardia. Stress-strain curves were calculated, filtered and resampled. The results show that Choi-Vito model was found to provide the best fit to the LV, the polynomial (Anisotropic) model to RV, the Four-Fiber Family model to RV, Holzapfel (2000) to RV, Holzapfel (2005) to RV and the Fung model to LV.

## 1. Introduction

The heart muscle is one of the most complex tissues in the animal body. It is heterogenous as well as anisotropic in nature [1-4]. By function, it acts under both active (during systole) and passive (in diastole) forces to pump blood throughout the body. The right side of the body handles oxygen-poor blood and pumps it to the lungs while the left side handles oxygen-rich blood and pumps it to the rest of the body. These powerful blood pumping functions are carried out by the walls called the ventricles. In between these walls is the separating wall called the intraventricular septum (IVS), which is quite dynamic during each cardiac cycle, contracting with the ventricles during systole and acting as a support structure against which the free wall of the right ventricle contracts. These differences in the cardiac muscle functions only make it obvious that these walls have different properties as functionality is mostly governed by property [5]. Golob et al. [5] reports that the many forms of heart disease are accompanied by a corresponding degradation in biomechanical performance of the ventricular myocardial tissue.

Therefore, the understanding of the mechanical properties of the soft tissue is critical in the development of reliable computational models [6-11]. The understanding ventricular mechanics can elucidate our understanding of its complex mechanical behaviour under different operating conditions and may enhance the understanding of heart function and failure [12-16]. This knowledge is essential to understanding the mechanisms of heart malfunctioning and is key to devising effective therapies [17-19]. However, most studies of the behaviour of heart muscles have focussed on the left side of the heart because of its relatively higher pumping demands than the right side [20-24]. The myocardial tissue plays a central role in blood pumping; therefore, it is the focus of the present study. Recent studies have shown that the right ventricle myocardium has its own unique properties that are distinct from those of the other myocardia. Sacks and Chuong [25] show that the right ventricle is different from the left ventricle in terms of fiber direction stiffnesses and degree of anisotropy. It is further shown that the lung-related diseases such as Covid-19 seem to affect the RV myocardial tissue more than the LV. Although the authors have studied the varying properties in the rat myocardia, it is necessary to investigate such differences in the myocardial tissue of bigger animals that are assumed to be closer to human heart anatomy than the smaller animals.

In this paper, the differences in myocardial tissue excised from the three main walls of the sheep heart are studied by examining the computational performances of six different constitutive models, namely, the Fung [26], Choi-Vito [27], Holzapfel (2000) [28], Holzapfel (2005) [29], Four-Fiber family [30, 31], and the Polynomial (Anisotropy) [32] hyperelastic models. The formalisms of these different models are examined and correlated with their computational performances in the three myocardial tissues. In the rat myocardia, both experimental and model investigations show that the RV is more anisotropic than the LV and MDW myocardial tissues in agreement with the findings of Sacks and Chuong [26]. The complex fiber-orientation and higher degree of anisotropy in the RV renders it harder to be accurately modelled by constitutive models that merely assume orthotropy. The rationale behind the investigation of these different hyperelastic material models is that it is relatively easier and less costly to study the biomechanics of the myocardial tissue on the mathematical models than real animal models. The experimental studies of real animal myocardial tissues pose huge limitations due to relatively higher costs and ethical demands than mathematical models. The testing procedures and sample preparation are highly demanding for real animal model testing. On the other hand, mathematical models do not require complicated specimen preparation once their parameters are accurately determined. In most cases, there are no demanding ethical clearance application processes. Furthermore, the mathematical models provide the researchers with opportunities for studying the effects of numerous varying physiological conditions without any adverse effects to life.

Li [33] recommended that there is need for more uniaxial and biaxial tensile testing in border and remote myocardial tissues in different time scales. It may be added that the testing must be extended to as many different loading cases as practically feasible in order to understand the mechanical behaviour of myocardial tissue. This is even more essential for the understanding of the healing process in the infarcted myocardium so that more effective therapeutic strategies may be devised that can arrest its progression to total heart failure.

## 2. MATERIALS AND METHODS

### a. Tissue preparation

Sheep hearts (N = 10) of unknown heart conditions and age were collected from the local abattoir one (1) hour after slaughtering. During collection, the hearts were place in the cooler box for delivering to the Biomechanics Lab and arrived in the lab after an hour drive. Immediately after arrival the sheep hearts were then prepared to ensure readiness for mechanical testing. All samples were then stored in a 0.9% NaCl physiological saline solution (PSS) for 30 minutes before preparation. After this, the hearts were then prepared by cutting square of 18 mmx 18 mm.

### b. Biaxial mechanical testing

The biaxial testing methodology and protocol was mainly adopted from the previously published work from the research group [9, 34]. Using the direction from the base to the apex and isolating papillary muscle as the main direction, the 18 mm x 18 mm square sample were cut from the LV, RV and IVS. In this work, the longitudinal direction which is along the papillary muscle was used with corresponding circumferential direction at 90° of the longitudinal direction. In this study the microstructural coordinate system (fibre and cross-fibre direction) was not used because the imaging to locate fiber direction was not performed. Instead, the longitudinal and circumferential system was considered. All heart wall samples in the LV, RV and IVS regions were clapped using BioTester 5000 CellScale, Waterloo, Canada apparatus. The square sample myocardial tissues were clamped at approximately 16 mm x 16 mm BioRake hooks. The preload of 0.5 mN was then applied in each sample during testing. To reduce the built-in stresses, all samples were subjected to a preconditioning as described in the previous study [34]. Each tissue as loaded to 0.4 strain equally in both longitudinal and circumferential direction. Toi mimic the body temperature, the 0.9% NaCl physiological saline solution (PSS) was then heated to 37°C before biaxial mechanical testing. The biaxial mechanical testing only begun after the saline solution reached a temperature of 37°C. The cross-sectional area of the samples was determined by measuring the thickness of the sample using Vernier calliper.

### c. Constitutive hyperelastic modelling

Typically two approaches are used for representation of anisotropy in biological soft tissues: the Green-Lagrange tensor-based approach [35, 17, 18, 19] and the strain invariants-based approach. The Green-Lagrange components-based approaches involve expressing the strain energy density functions *W* as summations of different contributions of the Green-Lagrange strain tensors *E*_*ij*_. Ateshian and Costa [36] state that this formulation allows the uncoupling of the strain energy function into dilatational and distortional parts which facilitates the computational implementation of incompressibility. However, it is reported by Chagnon et al [37] that these constitutive models have many material properties which affect their polyconvexity and make them relatively less stable. The other difficulty with these models is that the material parameters have no physical significance, hence making them very hard to fit. On the other hand, the strain invariant-based approaches express the strain energy density function *W* based on combinations of different isotropic and anisotropic functions. The isotropic and anisotropic functions are given as strain invariants, *I*_*k*_. There are three different implementations of this in the literature comprising polynomial, power, and exponential implementations, with the latter being the most popular because it includes the strain hardening effect in soft tissue. There are other forms of the strain invariant-based methods in the literature that implement logarithmic or tangent functions. However, these forms are specifically suitable for the modelling of moderate deformation especially before soft tissue activation. A significant characteristic of the strain invariant approaches is the inclusion of the anisotropic function of strain invariants *I*_4_ and/or *I*_6_ in their strain energy density functions. This study uses six anisotropic hyperelastic models two of which are Green-Lagrange tensor-based approaches and four are strain invariants-based approaches. The expressions for their strain energy density functions are given in **Table 1**.

**Table 1.**
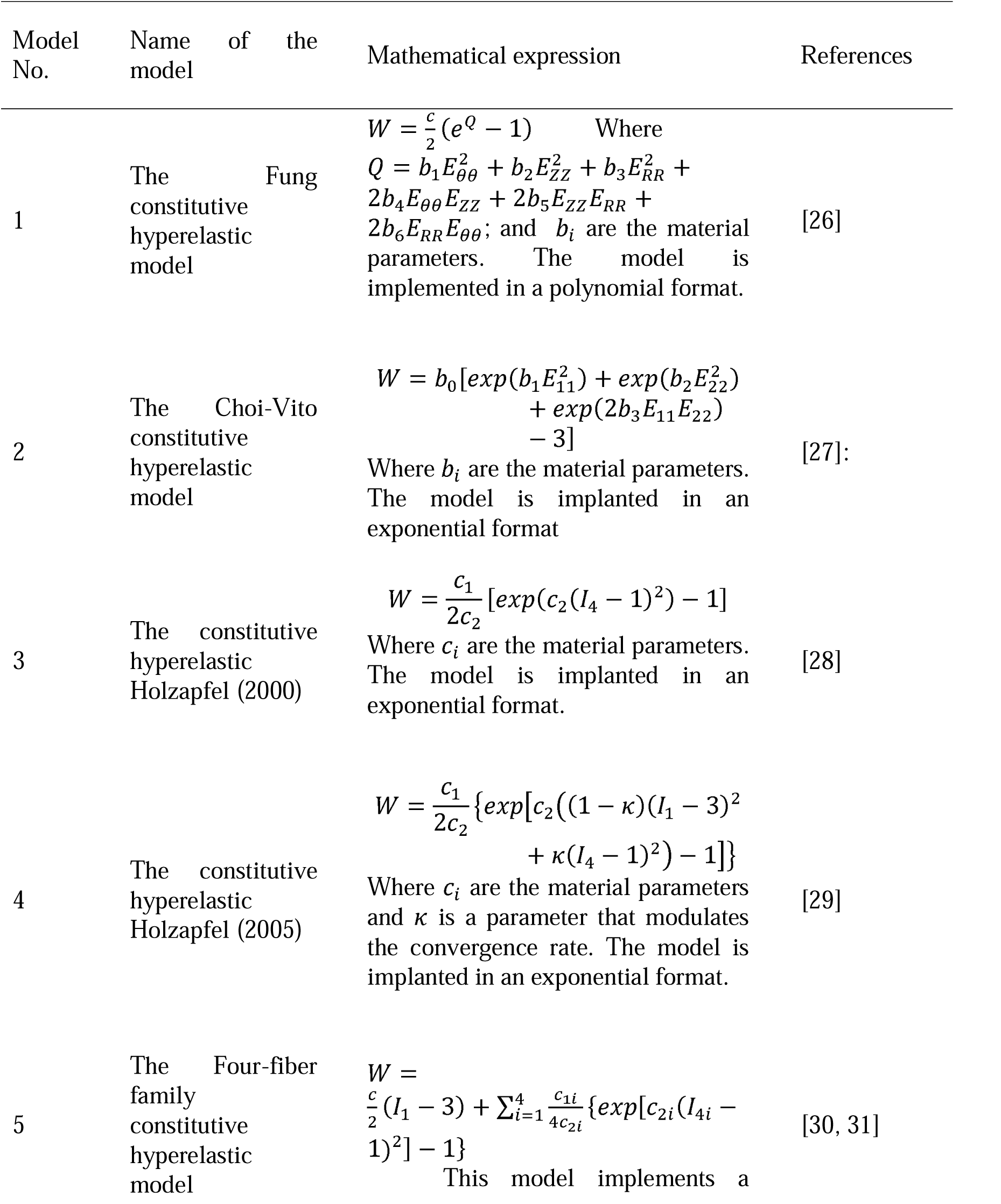
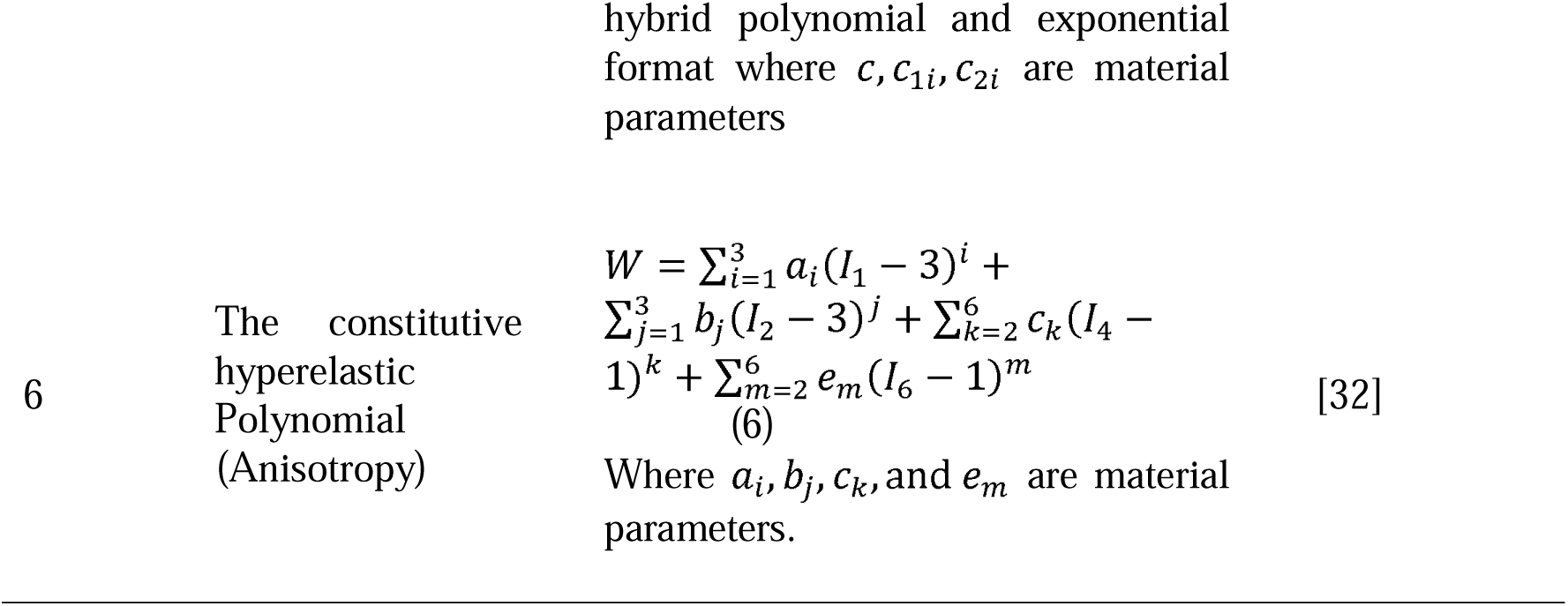
List of all hyperelastic constitutive models utilised in this study by fitting biaxial tensile data of the sheep different heart regions/walls (i.e. Left ventricle, Right ventricle, and Septal wall).

| Model No. | Name of the model | Mathematical expression | References |
| --- | --- | --- | --- |
| 1 | The constitutive hyperelastic model of Fung | $W = \frac{c}{2}(e^Q - 1) \quad \text{Where}$ $Q = b_1 E_{\theta\theta}^2 + b_2 E_{ZZ}^2 + b_3 E_{RR}^2 + 2b_4 E_{\theta\theta} E_{ZZ} + 2b_5 E_{ZZ} E_{RR} + 2b_6 E_{RR} E_{\theta\theta}; \text{ and } b_i \text{ are the material parameters. The model is implemented in a polynomial format.}$ | [26] |
| 2 | The Choi-Vito constitutive hyperelastic model | $W = b_0 [\exp(b_1 E_{11}^2) + \exp(b_2 E_{22}^2) + \exp(2b_3 E_{11} E_{22}) - 3]$<br>Where $b_i$ are the material parameters. The model is implanted in an exponential format | [27]: |
| 3 | The constitutive hyperelastic Holzapfel (2000) | $W = \frac{c_1}{2c_2} [\exp(c_2 (I_4 - 1)^2) - 1]$<br>Where $c_i$ are the material parameters. The model is implanted in an exponential format. | [28] |
| 4 | The constitutive hyperelastic Holzapfel (2005) | $W = \frac{c_1}{2c_2} \{ \exp[c_2 ((1 - \kappa)(I_1 - 3)^2 + \kappa(I_4 - 1)^2) - 1] \}$<br>Where $c_i$ are the material parameters and $\kappa$ is a parameter that modulates the convergence rate. The model is implanted in an exponential format. | [29] |
| 5 | The Four-fiber family constitutive hyperelastic model | $W = \frac{c}{2} (I_1 - 3) + \sum_{i=1}^4 \frac{c_{1i}}{4c_{2i}} \{ \exp[c_{2i} (I_{4i} - 1)^2] - 1 \}$<br>This model implements a | [30, 31] |

## 3. RESULTS

The thickness of the LV, RV and IVS were found to be 14.5 ± 0.85, 7.2 ± 0.25, 5.2 ± 0.13 mm, respectively. These thicknesses were used to calculate the true engineering stress and strain in both longitudinal and circumferential directions. To calculate the engineering stress, the cross-sectional area was calculated using the thickness and the breadth (16 ± 0.012 mm) of the tissue sample.

Tables 2-7 show the variations in both the fitting and material parameters of the six hyperelastic constitutive models. As expected, the variations in the material parameters over the ten tests are extremely large, which is indicative of the probabilistic nature of the material properties of the myocardium from one sample to another. It is therefore very hard to apply a set of material parameters obtained from one specimen to fit experimental data obtained from a different specimen, even for the same model. It is however very encouraging to see how small the variations are for the models’ fit parameters, (e.g. the coefficient of determination R^2^), for all the models under study.

**Table 2.** Biaxial mechanical data fitted on **Fung hyperelastic constitutive model** to determine the material parameters (c, b_1_, b_2_, b_3_, b_4_, b_5_ and b_6_). The average parameters were calculated from the material parameters of each tissue sample.

| Parameters | LV | IVS | RV |
| --- | --- | --- | --- |
| Coefficient of Determination ($R^2$) | $0.978 \pm 0.012$ | $0.921 \pm 0.122$ | $0.953 \pm 0.045$ |
| Correlation Coefficient (r) | $0.990 \pm 0.005$ | $0.961 \pm 0.070$ | $0.984 \pm 0.023$ |
| Normalised Error (NE) | $0.126 \pm 0.031$ | $0.162 \pm 0.159$ | $0.145 \pm 0.099$ |
| Norm. RMS Error (NRMSE) | $0.150 \pm 0.037$ | $0.201 \pm 0.177$ | $0.184 \pm 0.121$ |
| $c$ | $2.199 \pm 1.380$ | $35.178 \pm 47.660$ | $30.275 \pm 63.779$ |
| $b_1$ | $2.721 \pm 0.580$ | $0.301 \pm 0.307$ | $0.678 \pm 2.536$ |
| $b_2$ | $0.186 \pm 0.352$ | $0.126 \pm 0.471$ | $0.420 \pm 0.460$ |
| $b_3$ | $-1.098 \pm 0.509$ | $-0.070 \pm 0.447$ | $0.293 \pm 1.972$ |
| $b_4$ | $-0.531 \pm 0.307$ | $0.312 \pm 0.339$ | $0.283 \pm 1.847$ |
| $b_5$ | $1.331 \pm 0.928$ | $-0.247 \pm 0.358$ | $-1.204 \pm 2.281$ |
| $b_6$ | $-1.523 \pm 0.602$ | $-0.194 \pm 0.238$ | $-0.066 \pm 0.965$ |

**Table 3.**
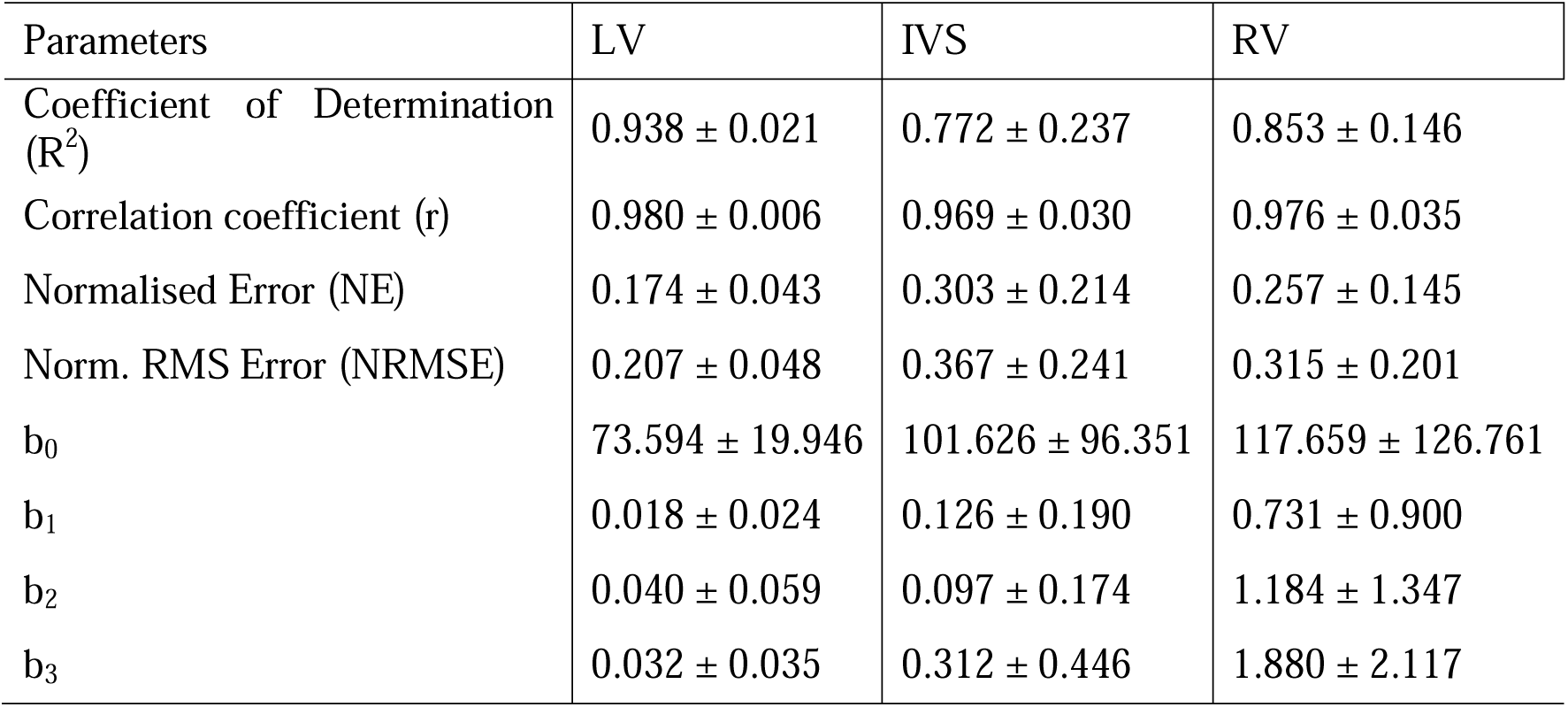
Biaxial mechanical data fitted on **Choi-Vito hyperelastic constitutive model** to determine the material parameters (b_0_, b_1_, b_2_ and b_3_). The average parameters were calculated from the material parameters of each tissue sample.

| Parameters | LV | IVS | RV |
| --- | --- | --- | --- |
| Coefficient of Determination ($R^2$) | $0.938 \pm 0.021$ | $0.772 \pm 0.237$ | $0.853 \pm 0.146$ |
| Correlation coefficient (r) | $0.980 \pm 0.006$ | $0.969 \pm 0.030$ | $0.976 \pm 0.035$ |
| Normalised Error (NE) | $0.174 \pm 0.043$ | $0.303 \pm 0.214$ | $0.257 \pm 0.145$ |
| Norm. RMS Error (NRMSE) | $0.207 \pm 0.048$ | $0.367 \pm 0.241$ | $0.315 \pm 0.201$ |
| $b_0$ | $73.594 \pm 19.946$ | $101.626 \pm 96.351$ | $117.659 \pm 126.761$ |
| $b_1$ | $0.018 \pm 0.024$ | $0.126 \pm 0.190$ | $0.731 \pm 0.900$ |
| $b_2$ | $0.040 \pm 0.059$ | $0.097 \pm 0.174$ | $1.184 \pm 1.347$ |
| $b_3$ | $0.032 \pm 0.035$ | $0.312 \pm 0.446$ | $1.880 \pm 2.117$ |

**Table 4.** Biaxial mechanical data fitted on **Polynomial (Anisotropic) hyperelastic constitutive model** to determine the material parameters (a_1_, a_2_, a_3_, b_1_, b_2_, b_3_, c_2_, c_3_, c_4_, c_5_, c_6_, φ). The average parameters were calculated from the material parameters of each tissue sample.

| Parameters | LV | IVS | RV |
| --- | --- | --- | --- |
| Coefficient of Determination ($R^2$) | $0.974 \pm 0.023$ | $0.948 \pm 0.060$ | $0.986 \pm 0.009$ |
| Correlation coefficient (r) | $0.117 \pm 0.043$ | $0.152 \pm 0.108$ | $0.107 \pm 0.045$ |
| Normalised Error (NE) | $0.989 \pm 0.009$ | $0.978 \pm 0.026$ | $0.994 \pm 0.004$ |
| Norm. RMS Error (NRMSE) | $0.097 \pm 0.038$ | $0.118 \pm 0.082$ | $0.084 \pm 0.033$ |
| $a_1$ | $0.787 \pm 0.397$ | $0.516 \pm 0.219$ | $0.701 \pm 0.268$ |
| $a_2$ | $0.496 \pm 0.681$ | $0.187 \pm 0.382$ | $0.670 \pm 0.506$ |
| $a_3$ | $0.044 \pm 0.296$ | $0.143 \pm 0.121$ | $0.333 \pm 0.288$ |
| $b_1$ | $0.019 \pm 0.050$ | $-0.019 \pm 0.031$ | $0.000 \pm 0.060$ |
| $b_2$ | $0.006 \pm 0.040$ | $0.008 \pm 0.027$ | $-0.003 \pm 0.043$ |
| $b_3$ | $0.027 \pm 0.090$ | $-0.019 \pm 0.060$ | $-0.024 \pm 0.034$ |
| $c_2$ | $-0.212 \pm 0.250$ | $0.051 \pm 0.198$ | $-0.122 \pm 0.072$ |
| $c_3$ | $-0.014 \pm 0.131$ | $0.171 \pm 0.205$ | $-2.007 \pm 5.971$ |
| $c_4$ | $0.177 \pm 0.140$ | $0.084 \pm 0.094$ | $0.034 \pm 0.063$ |
| $c_5$ | $0.006 \pm 0.099$ | $-0.072 \pm 0.270$ | $-0.030 \pm 0.108$ |
| $c_6$ | $-0.038 \pm 0.188$ | $-0.118 \pm 0.112$ | $0.029 \pm 0.061$ |
| **f** | $0.027 \pm 0.068$ | $-0.016 \pm 0.034$ | $-0.017 \pm 0.098$ |

**Table 5.** Biaxial mechanical data fitted on **Four-fiber family hyperelastic constitutive model** to determine the material parameters (c, c_11_, c_21_, c_12_, c_22_, c_134_, c_234_ and φ_0_). The average parameters were calculated from the material parameters of each tissue sample.

| Parameters | LV | IVS | RV |
| --- | --- | --- | --- |
| Coefficient of Determination ($R^2$) | $0.977 \pm 0.010$ | $0.943 \pm 0.052$ | $0.986 \pm 0.052$ |
| Correlation Coefficient (r) | $0.132 \pm 0.027$ | $0.147 \pm 0.093$ | $0.084 \pm 0.093$ |
| Normalised Error (NE) | $0.16 \pm 0.035$ | $0.178 \pm 0.108$ | $0.105 \pm 0.108$ |
| Norm. RMS Error (NRMSE) | $0.99 \pm 0.005$ | $0.975 \pm 0.024$ | $0.994 \pm 0.024$ |
| $c$ | $2.075 \pm 0.832$ | $0.328 \pm 0.265$ | $0.211 \pm 0.265$ |
| $c_{11}$ | $97.187 \pm 304.567$ | $0.665 \pm 0.966$ | $1.172 \pm 0.966$ |
| $c_{21}$ | $0.924 \pm 0.454$ | $1.357 \pm 0.648$ | $1.030 \pm 0.648$ |
| $c_{12}$ | $2.357 \pm 1.434$ | $1.98 \pm 1.317$ | $0.731 \pm 1.317$ |
| $c_{22}$ | $1.072 \pm 0.324$ | $0.539 \pm 0.574$ | $1.139 \pm 0.574$ |
| $c_{134}$ | $2.132 \pm 0.782$ | $0.792 \pm 0.670$ | $1.599 \pm 0.670$ |
| $c_{234}$ | $1.765 \pm 0.623$ | $0.908 \pm 1.084$ | $0.974 \pm 1.084$ |
| $\phi_0$ | $0.85 \pm 0.560$ | $0.981 \pm 0.723$ | $0.895 \pm 0.723$ |

**Table 6.** Biaxial mechanical data fitted on **Holzapfel (2000) hyperelastic constitutive model** to determine the material parameters (μ, k_1_, k_2_ and φ). The average parameters were calculated from the material parameters of each tissue sample.

| Parameters | LV | IVS | RV |
| --- | --- | --- | --- |
| Coefficient of Determination ($R^2$) | $0.96 \pm 0.024$ | $0.068 \pm 0.068$ | $0.984 \pm 0.014$ |
| Correlation Coefficient (r) | $0.129 \pm 0.037$ | $0.116 \pm 0.116$ | $0.089 \pm 0.031$ |
| Normalised Error (NE) | $0.156 \pm 0.046$ | $0.127 \pm 0.127$ | $0.112 \pm 0.039$ |
| Norm. RMS Error (NRMSE) | $0.985 \pm 0.008$ | $0.033 \pm 0.033$ | $0.994 \pm 0.004$ |
| $\mu$ | $0 \pm 0.000$ | $0.015 \pm 0.015$ | $0.002 \pm 0.007$ |
| $k_1$ | $1.501 \pm 0.554$ | $0.408 \pm 0.408$ | $1.429 \pm 0.691$ |
| $k_2$ | $0.397 \pm 0.340$ | $0.258 \pm 0.258$ | $0.874 \pm 0.533$ |
| $\phi$ | $0.912 \pm 0.434$ | $0.389 \pm 0.389$ | $0.889 \pm 0.082$ |

**Table 7.** Biaxial mechanical data fitted on **Holzapfel (2005) hyperelastic constitutive model** to determine the material parameters (μ, k_1_, k_2_, φ, and ρ). The average parameters were calculated from the material parameters of each tissue sample.

| Parameters | LV | IVS | RV |
| --- | --- | --- | --- |
| Coefficient of Determination ($R^2$) | 0.966±0.022 | 0.953±0.057 | 0.986±0.013 |
| Correlation Coefficient (r) | 0.115±0.040 | 0.121±0.102 | 0.081±0.029 |
| Normalised Error (NE) | 0.142±0.050 | 0.156±0.117 | 0.102±0.038 |
| Norm. RMS Error (NRMSE) | 0.987±0.008 | 0.978±0.028 | 0.994±0.004 |
| $\mu$ | 0.032±0.054 | 0.073±0.131 | 0.109±0.140 |
| $k_1$ | 1.412±0.491 | 1.216±0.455 | 1.295±0.650 |
| $k_2$ | 0.226±0.278 | 0.1±0.206 | 0.552±0.665 |
| $\phi$ | 0.83±0.814 | 0.681±0.481 | 1.274±0.351 |
| r | 0.756±0.248 | 0.725±0.162 | 0.611±0.348 |

Figure 2 show the variations in Coefficient of Determination (R^2^) of all hyperplastic constitutive models across all samples.

**Figure 1.**
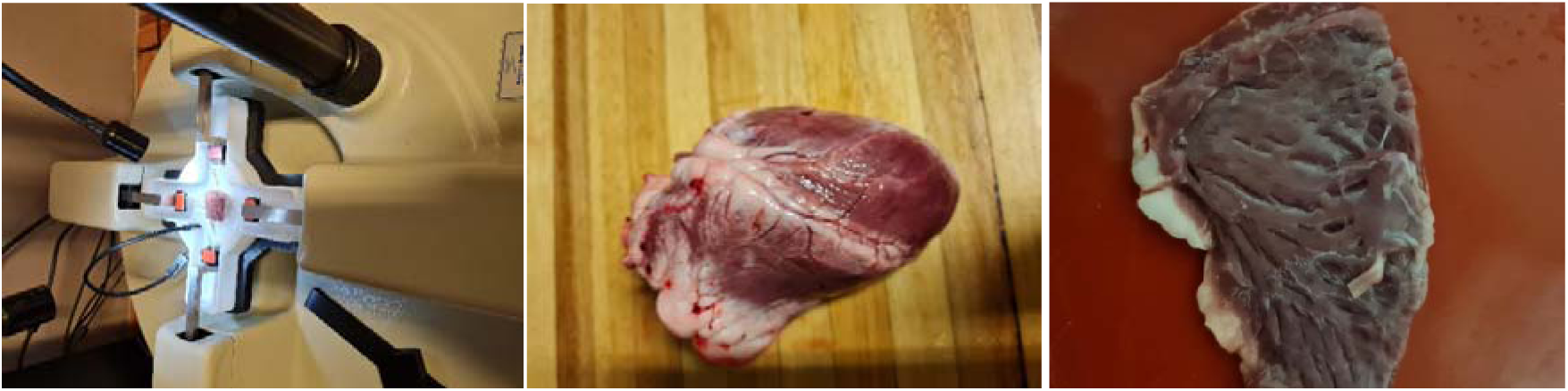
Experimental set up of equi-biaxial mechanical testing of sheep heart looking at different three regions including Left ventricle (LV), septal wall (IVS) and Right ventricle (RV)

**Figure 2.**
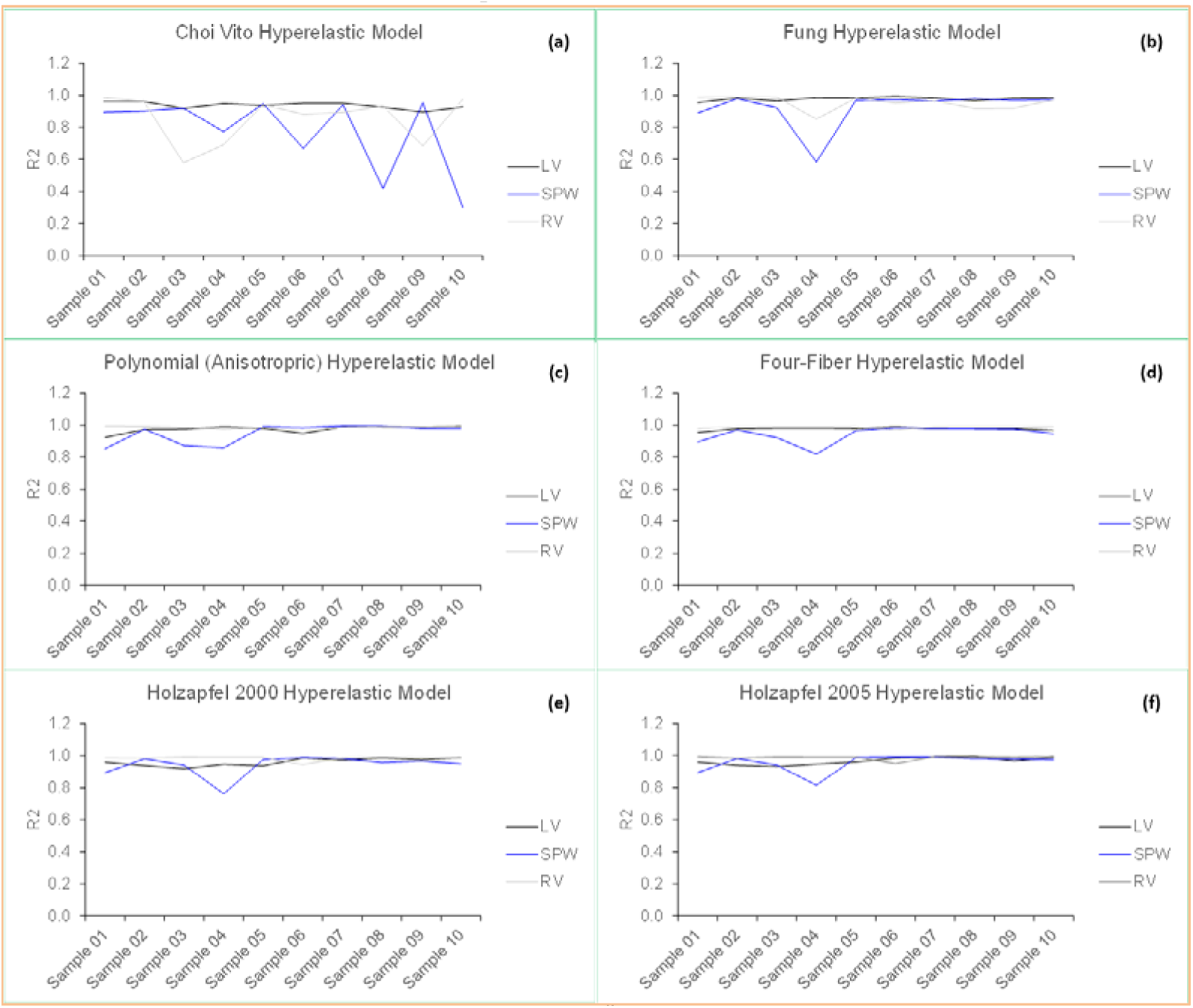
Coefficient of Determination (R^2^) of all hyperplastic constitutive models across all samples considered during biaxial testing of sheep heart tissue in the regions of LV, SPQ and RV.

Figure 3 shows the Evaluation Index (EI) determined from the actual coefficient of determination (R^2^) of Fung, Polynomial (Anisotropic), Holzapfel 2000, Holzapfel 2005, Four-Fiber Family and Choi-Vito hyperelastic models across all the samples.

**Figure 3.**
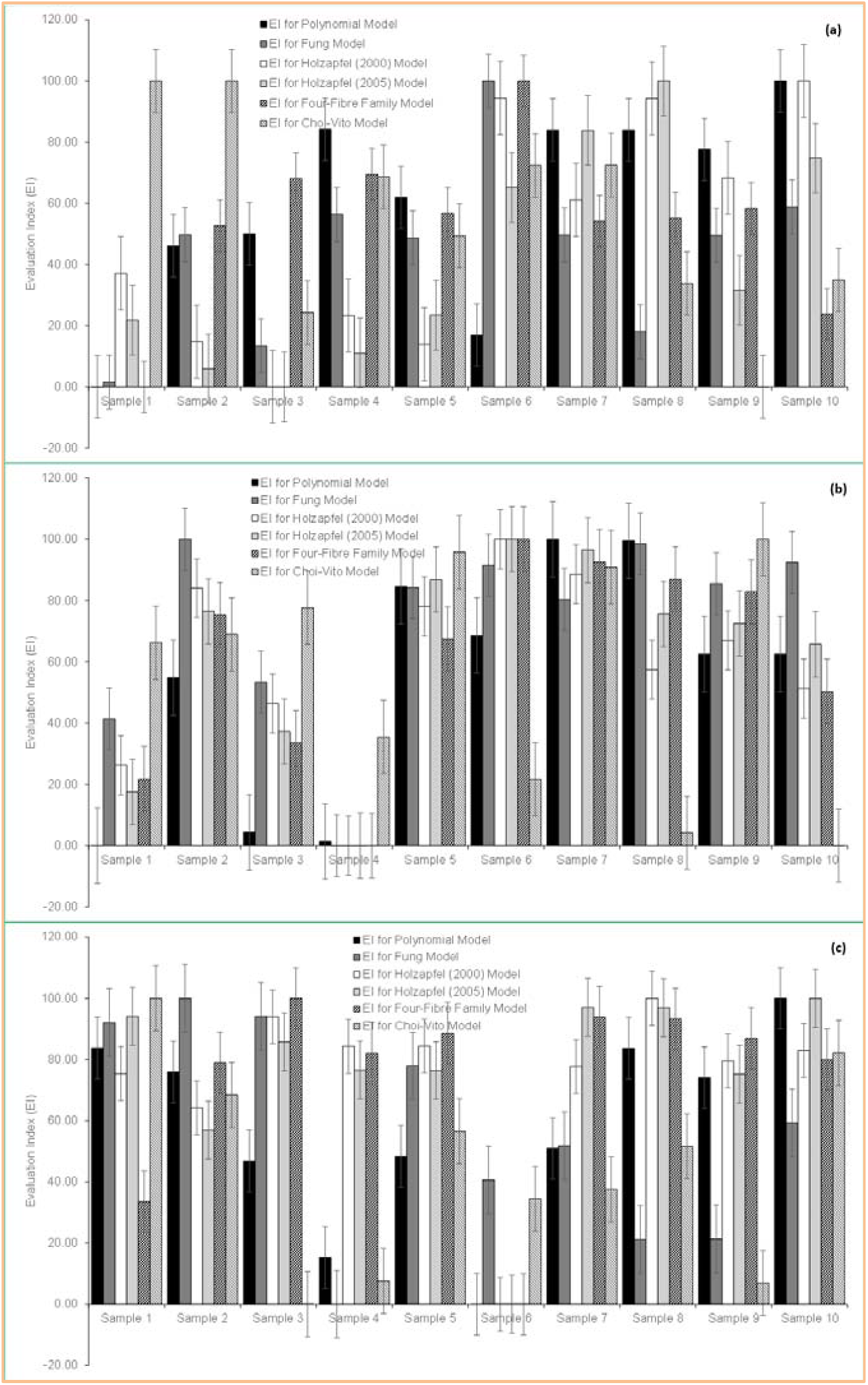
Evaluation Index (EI) determined from the actual coefficient of determination (R2) of Fung, Polynomial (Anisotropic), Holzapfel 2000, Holzapfel 2005, Four-Fiber Family and Choi-Vito hyperelastic models

## 4. DISCUSSION

The fitting performances of six different constitutive models on the different regions of the heart regions (LV, IVS and RV) were evaluated successfully. A direct comparison of hyperelastic constitutive models was made based on correlation coefficient (R^2^) and evaluation index (EI) for the LV, IVS and RV.

The R^2^ coefficients of average curves of LV, IVS and RV for six different constitutive models are presented in Table 8. In summary, it was found that Choi-Vito model was found to provide the best fit to the LV, the polynomial (Anisotropic) model to RV, the Four-Fiber Family model to RV, Holzapfel 2000 to RV, Holzapfel 2005 to RV and the Fung model to LV. The R squared (R^2^) was utilised as a deciding factor, however it is reported that averaging the stress-strain curve is best option [1]. It has been observed that the average R Squared (R^2^) of experimental fitted data may not reproduce average hyperelatic constitutive model. Also, the individual R squared (R^2^) generated from each sample (from N = 10) fitted in six hyperelastic models are compared. This approach may assist in coming to the right conclusion since the plotted trend of selected constitutive models for each R^2^.

**Table 8.** R^2^ values for all heart wall regions (i.e. LV, RV and IVS)

| Model | LV | IVS | RV |
| --- | --- | --- | --- |
| Fung Model | 0.98 | 0.92 | 0.95 |
| Polynomial (Anisotropic) | 0.97 | 0.95 | 0.99 |
| Holzapfel (2000) | 0.96 | 0.94 | 0.98 |
| Holzapfel (2005) | 0.97 | 0.95 | 0.99 |
| Four-Fiber Family | 0.98 | 0.94 | 0.99 |
| Choi-Vito | 0.94 | 0.77 | 0.85 |

**Table 9.** P-values for R^2^: Cross-wall variation of all heart wall regions (i.e. LV, RV and IVS)

| | P-values for $R^2$: Cross-wall variation in a particular direction | | |
| --- | --- | --- | --- |
|  | LV - IVS | LV - RV | IVS - RV |
| **Choi- Vito** | 0.041 | 0.085 | 0.370 |
| **Polynomial (Anisotropic)** | 0.213 | 0.152 | 0.067 |
| **Four-Fiber Family** | 0.063 | 0.014 | 0.019 |
| **Holzapfel 2000 Model** | 0.376 | 0.017 | 0.061 |
| **Holzapfel 2005 Model** | 0.527 | 0.024 | 0.095 |
| **Fung Model** | 0.155 | 0.1066 | 0.436 |

Figure 4 shows the box plot of the six different constitutive hyperelastic model in comparing the R^2^ of the LV, IVS and RV regions of the sheep heart.

**Figure 4.**
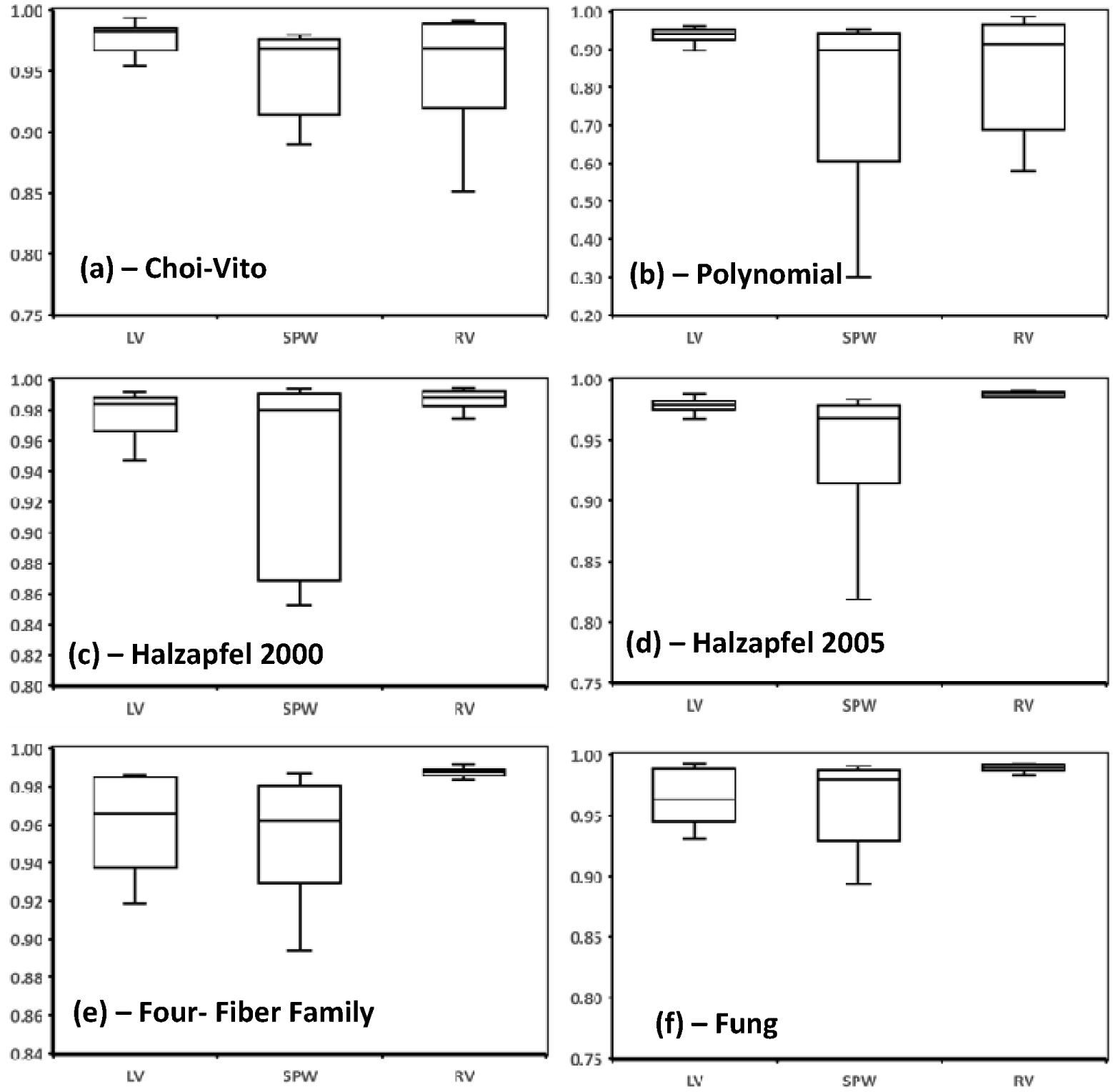
Box plot of (a) and Choi-Vito, (b) Polynomial, (c) Halzapfel (2000), (d) Halzapfel (2005), (e) Four fiber and (f) Fung model for (R^2^) of LV, IVS and RV Using the R^2^, it must also be noted that there was significant cross-directional variances in the LV - RV, lightly significant in IVS - RV and non-significant in the LV-IVS as shown in **Table 9**.

For the Fung model, the differences in R^2^ between the LV - IVS were found to be significant (p = 0.155) and between LV-RV were also insignificant (p=0.107). Additionally, no noteworthy differences in R^2^ were found between the IVS -RV (p = 0.436).

For the Choi-Vito model, the differences in R^2^ between the LV - IVS were found to be significant (p = 0.041) and between LV - RV were also insignificant (p=0.085). Additionally, no noteworthy differences in R2 were found between the IVS - RV (p = 0.370).

For the Polynomial (Anisotropic) model, the differences in R2 between the LV - IVS were found to be insignificant (p = 0.213) and between LV-RV were also insignificant (p=0.152). Additionally, no noteworthy differences in R^2^ were found between the IVS -RV (p = 0.067). For the Four-Fiber Family model, the differences in R^2^ between the LV-IVS were found to be less significant (p = 0.063). There was significant differences between LV-RV (p = 0.014) and also IVS - RV (p = 0.019).

For the Holzapfel 2000 model, the differences in R^2^ between the LV-IVS were found to be insignificant (p = 0.376), yet significant between LV-RV were (p=0.017). Additionally, less differences in R^2^ were found between the IVS - RV (p = 0.061).

The Holzapfel 2005, the differences in R^2^ between the LV - IVS were found to be insignificant (p = 0.527) yet significant between LV-RV were (p=0.024). Additionally, less differences in R^2^ were found between the IVS-RV (p = 0.095).

For the Fung model, differences in R^2^ between the LV – IVS, LV - RV and IVS - RV were found to be insignificant (p = 0.155, 0.1066, 0.436, respectively).

For the LV, the Fung model was found to have EI of 100. This means that it is the best performance constitutive model when compared to other five hyperelastic model considered in this study. The Holzapfel 2005 constitutive model was found to have the EI of 100 and therefore regarded best performances when fitted on IVS of the sheep heart. Finally, for the RV, the four fibre-family was the best performer with EI of 100. The Choi-Vito model had the worst performance in the LV, IVS and RV region, which shows myocardial tissue behaviour cannot be accurately represented by model that do not incorporate fiber orientation and anisotropy.

**Figure 9.**
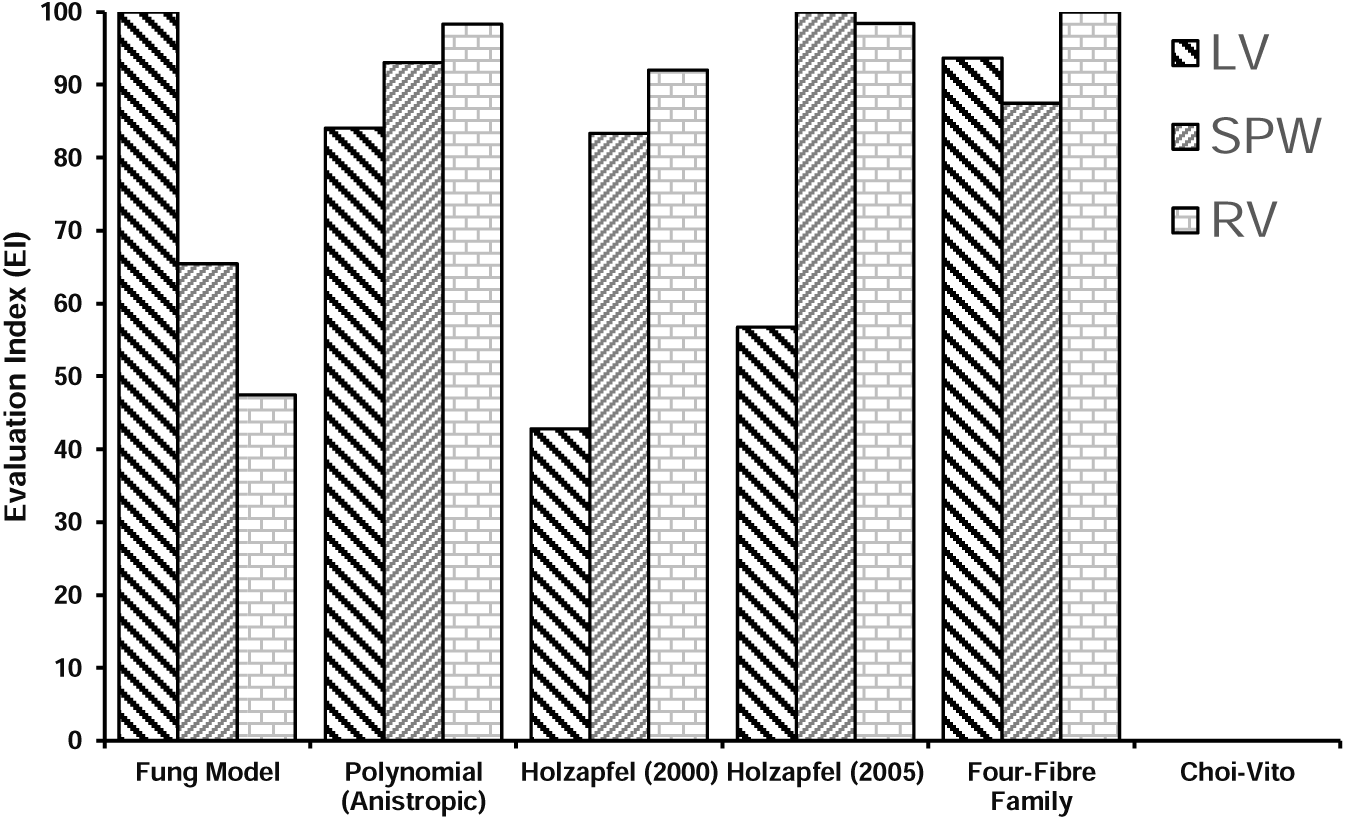
Evaluation Index (EI) calculated from the average R^2^ of Fung, Polynomial (Anisotropic), Holzapfel 2000, Holzapfel 2005, Four-Fiber Family and Choi-Vito hyperelastic models

## 5. CONCLUSION

The present study is a crucial step towards more robust constitutive modelling of healthy myocardium by the characterisation of the mechanical properties of various tissues under biaxial loading conditions. Biaxial mechanical tests results conducted on the sheep heart fibre tissue from the LV, IVS and RV were fitted to six hyperelastic models.

The present study demonstrated the importance of fitting correct models for the different regions of the sheep heart to get more accurate material properties. This is crucial as these material characteristics can be used for more accurate computer simulations of the cardiac mechanical function. These material parameters could be confidently utilised in the future for development Finite Element Models (FEM) in understanding how myocardial infarction affect the global functioning of the heart. Good mechanical properties of the heart region can lead to improved interventions and therapies from knowledge gained from FEM models.

## Acknowledgements

This study was supported financially by the University of South Africa. Any opinion, findings and conclusions or recommendations expressed in this publication are those of the authors and therefore the University of South Africa does not accept any liability in regard thereto.

## Author contributions

F.N, and T.P contributed to conception, design, data acquisition, analysis, and interpretation, drafted the manuscript, H.N, T.P and F.N contributed to the interpretation of the data and critically revised the manuscript, and finally F.N, H.N, and T.P contributed to conception, design, data analysis, and interpretation, drafted and critically revised the manuscript. All authors gave final approval and agree to be accountable for all aspects of the work.

## Corresponding author

Correspondence to Fulufhelo Nemavhola

## Declarations Ethics approval and consent to participate

The animal study ethics was approved by the College of Science Engineering and technology Ethic Committee of the University of South Africa on 4^th^ of September 2021 under reference number 2019/CSET_SOE/ARN/001

## Consent for publication

Not applicable.

